# Channel Capacity for Time-Resolved Effective Connectivity in Functional Neuroimaging

**DOI:** 10.64898/2026.03.28.714906

**Authors:** Jianan Jian, Benjamin Li, Nurahmed Multezem, Francesca Mandino, Evelyn Lake, Nan Xu

## Abstract

Understanding how brain regions influence one another over time is a central goal of neuroscience. However, existing approaches to effective connectivity often involve tradeoffs among mechanistic interpretability, computational scalability, and time-resolved estimation. Here, we introduce information channel capacity, a model-based measure of directed information transfer between brain regions, and combine it with a sliding-window framework to estimate time-varying directional interactions. We validate channel capacity across multimodal neuroimaging datasets in humans and rodents because this breadth is needed to test three complementary properties that no single dataset can establish alone: sensitivity to evoked information transfer, specificity against false-positive directional effects, and the ability to capture meaningful temporal variability in directed brain-network interactions. Human motor-task fMRI tests sensitivity, showing that channel capacity detects task-related increases in directed interactions and stronger directional effects during task than during rest in motor-related regions. Concurrent rat local field potential (LFP)-fMRI tests specificity, showing minimal spurious directional asymmetry relative to scan-to-scan variability and consistent temporal dynamics across neural and BOLD signals. Mouse Ca^2+^-fMRI tests temporal variability, showing that channel-capacity patterns identify reproducible connectivity states and transitions over time. Together, these results establish channel capacity as a physiologically grounded framework for measuring dynamic directional interactions across species and neuroimaging modalities.

## 1. Introduction

Coordinated interactions among distributed neural circuits give rise to perception, behavior, and higher-order cognition (Hutchison et al., 2013; Xu et al., 2021). Understanding how distributed brain regions communicate within and across these networks is therefore fundamental to explaining brain function. Functional neuroimaging techniques, particularly functional magnetic resonance imaging (fMRI), have made it possible to study these large-scale interactions in vivo by measuring dynamic signals across the whole brain (Friston, 2011; Ma et al., 2021; Mitra et al., 2015). At the same time, multimodal neuroimaging in rodents now makes it possible to relate whole-brain hemodynamic signals to more direct neural measurements, providing an opportunity to link large-scale functional organization with underlying neural activity (Lake et al., 2020; Mandino et al., 2025; Mitra et al., 2018; Thompson et al., 2013a; Zhang et al., 2020).

Functional connectivity (FC) analyses leverage these data to quantify statistical dependencies (i.e., correlations) between regions, enabling sensitive detection of the brain’s modular organization (Power et al., 2011; Thomas Yeo et al., 2011). Dynamic FC methods, such as sliding-window correlation (SWC) (Ma et al., 2021; Shakil et al., 2016), incorporate temporal variability of connectivity patterns and have successfully identified biomarkers across a range of neurological and psychiatric conditions (Damaraju et al., 2014; Vergara et al., 2018). Despite these advances, such approaches remain correlational in nature and are therefore limited in their ability to characterize directional neural interactions. This motivates the development of measures that can capture not only whether brain regions fluctuate together, but also how they influence one another over time.

Effective connectivity (EC) methods were developed to address this limitation (Friston, 2011). EC approaches span model-based frameworks (e.g., structural equation modeling and dynamic causal modeling) and predictive time-series formulations (e.g., Granger-causal analyses), among others, providing principled tools for characterizing directed interactions under different assumptions (Büchel & Friston, 1997; Friston et al., 2003; Seth et al., 2015; Smith et al., 2011). In practice, routine application at whole-brain scale can be challenging due to substantial modeling choices and computational considerations (Smith et al., 2011). Moreover, many EC formulations do not readily extend to time-varying settings, creating a practical tradeoff among mechanistic interpretability, computational scalability, and time-resolved estimation.

With the goal of developing a connectivity measure that is both practical and sensitive to temporal dynamics and directionality, our previous work proposed the prediction-correlation (p-corr) framework based on a causal model (Xu et al., 2017, 2021) and proposed a two-fold directed connectivity measure characterizing both the strength of directed FC and the duration of information transfer. By retaining the sensitivity of FC for detecting modular functional network organizations while introducing directionality through model-based prediction, p-corr was shown to be reliable and scalable for large-scale whole-brain fMRI analyses (Xu et al., 2017, 2021). We further extended p-corr to capture temporal dynamics using sliding-window techniques, namely sliding-window p-corr (SWpC), demonstrating reliable and sensitive dynamic directed connectivity in multimodal functional recordings, including fMRI and local field potentials (LFPs) (Xu et al., 2025, 2026).

Building on this framework, this paper introduces channel capacity as a novel connectivity measure that is dynamic, directional, reliable, and sensitive, while offering an information-theoretic characterization of neural communication. Channel capacity is a fundamental concept in information theory and communication engineering that quantifies the maximum rate at which information can be reliably transmitted through a communication channel under specified signal and noise constraints. Here, within the same directed modeling framework used to characterize interregional interactions, we estimate how much reliable directed information transfer is supported between brain regions under the fitted communication model. Importantly, channel capacity differs from many existing directed connectivity models in what it quantifies: rather than reporting a coupling parameter or a predictability statistic, e.g., (Büchel & Friston, 1997; Friston et al., 2003; Seth et al., 2015), it yields an information-theoretic measure with a direct interpretation as the maximum reliable directional information transfer supported by the fitted communication model. This is especially attractive for functional neuroimaging, where signal-to-noise ratio is often low and spatially heterogeneous, because channel-capacity estimation explicitly incorporates noise statistics.

Because no single dataset can establish all desired properties of a new directed-connectivity measure, we validated channel capacity across complementary human and rodent neuroimaging datasets. Human fMRI provides a clinically relevant noninvasive setting for studying large-scale directed interactions, whereas rodent multimodal datasets provide more direct neural measurements that help ground and validate these inferences. Using this cross-species framework, we show that channel capacity is sensitive to expected task-evoked directional effects in human motor-task fMRI, specific against spurious asymmetry in rat resting-state LFP-fMRI, and capable of capturing meaningful temporal variability and cross-modal correspondence in mouse Ca^2+^-fMRI. We further examine its relationship to correlation-based and predictive connectivity measures. Together, these analyses establish channel capacity as a physiologically grounded and computationally practical framework for studying dynamic directional interactions across species and neuroimaging modalities.

## 2. Methods

### 2.1. Modeling and Estimation Framework for Channel Capacity

#### 2.1.1. Channel model of effective connectivity

Effective connectivity from one ROI to another can be modeled as a communication channel. Over short timescales, this channel can be approximated by a linear time-invariant system (LTI) with a finite impulse response:

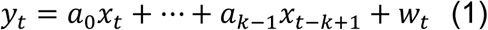

where 𝑥_𝑡_ is the signal in the sender ROI, 𝑦_𝑡_ is the signal in the receiver ROI, 𝑤_𝑡_ ∼ 𝒩(𝜇, 𝜎^2^) is an additive white Gaussian noise (AWGN) independent of 𝑥_𝑡_, and (𝑎_0_, …, 𝑎_𝑘−1_) are 𝑘 coefficients of impulse responses. Because negative correlations in GSR-processed data are difficult to interpret (Murphy et al., 2009; Saad et al., 2012), we restrict our analysis to nonnegative connectivity by imposing positivity constraints on these coefficients.

The frequency response of the system (1) is given by the discrete-time Fourier transform 𝐻(𝑓) = ∑^𝑘−1^ 𝑎_𝑗_𝑒^−2𝜋𝑖𝑗𝑓^. The equivalent input noise (EIN) has a power spectrum density (PSD) given by 𝑆_EIN_(𝑓) = 𝜎^2^/|𝐻(𝑓)|^2^. The channel capacity of this model is given by the water-filling solution (Gallager, 1968).:

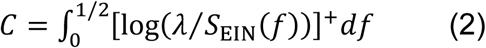

where [𝑐]^+^ ≔ max{0, 𝑐} denotes the rectifier function, and 𝜆 is a constant (“water level”) determined by the power constraint we impose on the sender signal:

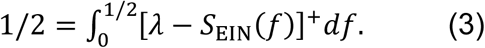

#### 2.1.2. Sliding-window framework of time-varying connectivity

For long-duration recordings, the temporal fluctuations in the system are no longer negligible. To capture time-varying connectivity, we adapt the system (1) into the sliding-window framework, by dividing the timeseries into 𝑀 temporal windows {𝑊_𝑖_}^𝑀^ and estimating the model parameters and connectivity measures on each of the windows (Fig. 1). Window length (denoted by 𝑛) and step size are meta-parameters selected based on established neurophysiological timescales of brain dynamics and the goals of the analysis.

**Fig. 1.**
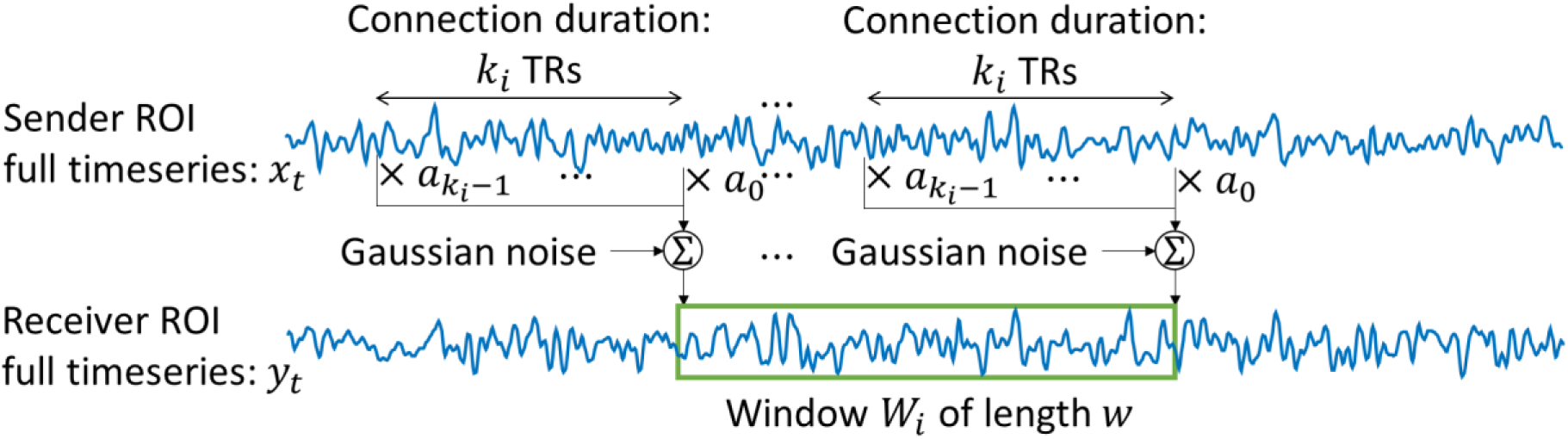
The sliding-window method applied with windows of size. 𝒏. The parameters of the system (1) are estimated on each window using 𝑛 observations of 𝑦_𝑡_ and their corresponding segments of 𝑥_𝑡_.

For each directed ROI pair within each window, the system (1) is rewritten as a linear regression matrix form:

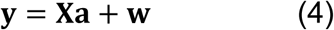

where 𝐲 is the column vector of 𝑛 samples of the receiver signal in the window, 𝐗 is the convolution matrix of size 𝑛-by-𝑘_𝑖_ constructed from the corresponding segments of the sender signal, 𝐚 is the column vector of 𝑘_𝑖_ unknown coefficients to be determined, and 𝐰 is the column vector of unknown noise. The model order 𝑘_𝑖_ in each window is a variable parameter selected by an information criterion (e.g., AIC/BIC) (Akaike, 1970, 1974; Schwarz, 1978; Sugiura, 1978). The connection duration within that window is defined as 𝑘_𝑖_TRs, representing the effective channel memory.

The vector 𝐲 and each column of 𝐗 are standardized by removing their means and dividing by their root mean squares (RMSs), so that the noise 𝐰 has zero-mean, and the power constraint imposed on the channel is enforced.

The coefficients 𝐚 within each window are estimated using the maximum likelihood estimator (MLE)

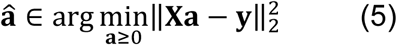

which can be obtained through a standard numerical solver (such as lsqnonneg in MATLAB). The MLE of the noise power 𝜎^2^ within each window is given by the mean square error (MSE) of the residuals: 𝜎̭^2^ = ‖𝐗𝐚̭ − 𝐲‖^2^⁄𝑛.

The same sliding-window framework can be applied to correlation-based connectivity measures. In each sliding-window, we computed the correlation between 𝐱 and 𝐲, called sliding-window correlation (SWC), as a non-directed connectivity measure for basic comparison. We also computed the correlation between the observed receiver signal 𝐲 and the model-predicted receiver signal 𝐲̭ = 𝐗𝐚̭, called sliding-window prediction-correlation (SWpC), as a directed connectivity measure for further comparison.

#### 2.1.3. Numerical estimation of channel capacity

After estimating the model parameters within each window, we estimate the channel capacity by approximating the water-filling solution. We approximate the discrete-time Fourier transform 𝐻(𝑓) by a discrete Fourier transform (DFT) on an equally spaced frequency grid 𝑓_𝑙_ = 𝑙⁄(𝑁 TR) for 𝑙 = 0, …, 𝑁 − 1, where TR denotes the repetition time and varies across imaging modalities. This is computed with a fast Fourier transform (FFT) of the vector 𝐚̭ zero-padded to length 𝑁^1^:

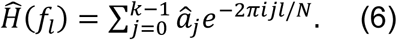

The values of 𝑆̭_EIN_(𝑓_𝑙_) = 𝜎̭^2^/𝐻̭(𝑓_𝑙_) are sorted in ascending order to form the sequence 𝑆̭^inc^ (𝑙). Starting from 𝑚 = 1, the candidate water level is computed as

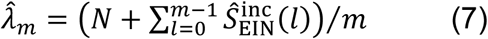

Iterate over 𝑚 until either 𝑚 = 𝑁 or 𝜆̭_𝑚_ ≤ 𝑆̭^inc^ (𝑚 + 1), then we obtain an estimation of the channel capacity

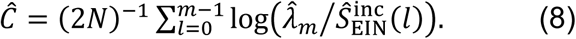

We repeat this calculation over all windows to obtain a time-resolved series of channel capacity aligned with the sliding-window scheme used for SWC and SWpC.

### 2.2. Data overview, acquisition, and preprocessing

We used three complementary neuroimaging datasets because no single dataset can establish all desired properties of the proposed measure. Human motor-task fMRI was used to test sensitivity to task-evoked directed interactions; rat resting-state LFP-fMRI was used to test specificity against spurious directionality and neural–BOLD correspondence; and mouse resting-state WF-Ca^2+^-fMRI was used to evaluate brain-wide temporal variability and cross-modal correspondence (see Table 1).

**Table 1.**
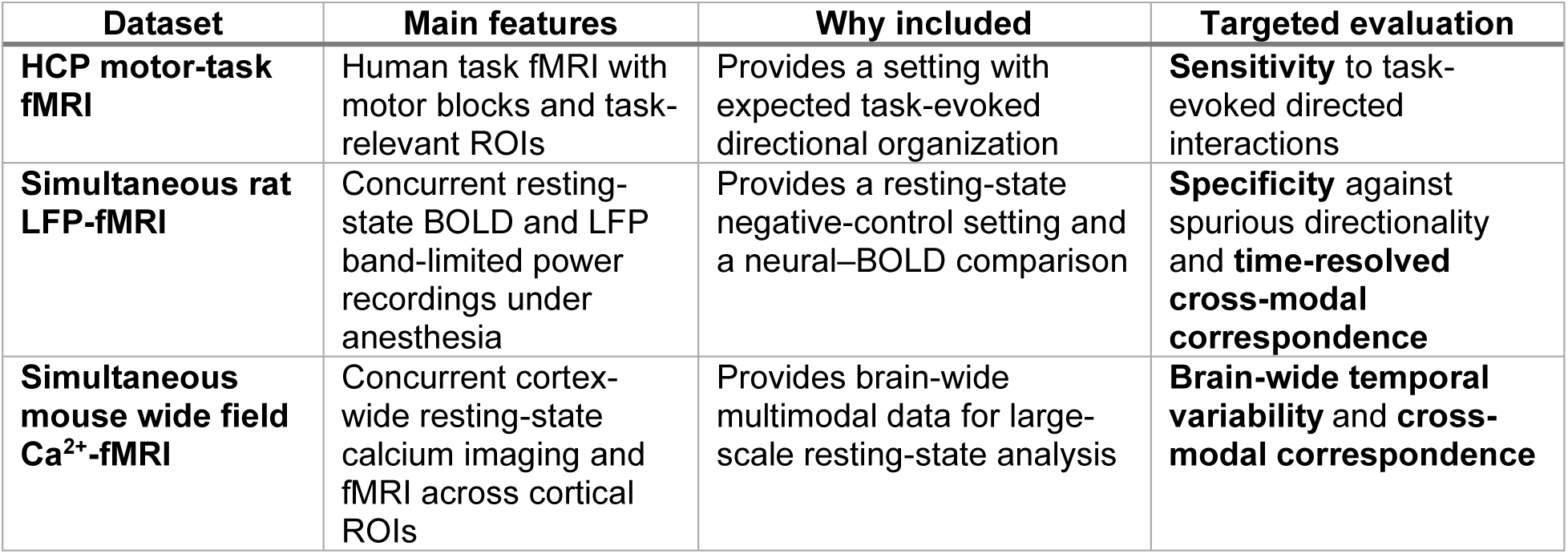
Overview of the datasets and their targeted roles in evaluating channel capacity.

| Dataset | Main features | Why included | Targeted evaluation |
| --- | --- | --- | --- |
| **HCP motor-task fMRI** | Human task fMRI with motor blocks and task-relevant ROIs | Provides a setting with expected task-evoked directional organization | **Sensitivity** to task-evoked directed interactions |
| **Simultaneous rat LFP-fMRI** | Concurrent resting-state BOLD and LFP band-limited power recordings under anesthesia | Provides a resting-state negative-control setting and a neural-BOLD comparison | **Specificity** against spurious directionality and **time-resolved cross-modal correspondence** |
| **Simultaneous mouse wide field Ca<sup>2+</sup>-fMRI** | Concurrent cortex-wide resting-state calcium imaging and fMRI across cortical ROIs | Provides brain-wide multimodal data for large-scale resting-state analysis | **Brain-wide temporal variability** and **cross-modal correspondence** |

#### 2.2.1. Motor-task fMRI data in humans

To test the sensitivity of channel capacity to task-evoked directed interactions, we analyzed motor-task fMRI from the Human Connectome Project (HCP) test–retest cohort by comparing channel capacity among task-specific ROIs during task and rest periods. These motor tasks are known to reliably activate the motor and somatosensory cortices, as well as the cerebellum (Barch et al., 2013).

The cohort included 45 participants (31 female; age range, 22–35 years). Each participant completed two motor-task scans with right-to-left and left-to-right phase encoding. Each scan consisted of thirteen 15-s blocks: an initial fixation block, followed by two repetitions of five consecutive movement blocks, each repetition followed by a fixation block of approximately 15 s. Each movement block included a 3-s cue and 12 s of movement (left foot, right foot, left hand, right hand, or tongue). Accordingly, each movement type was performed four times across the two scans per participant. The fixation blocks following each set of movement blocks served as the baseline condition for comparison with the movement blocks (see the block design in Fig. S1 of (Xu et al., 2026)).

Whole-brain fMRI data were downloaded from the HCP database as minimally preprocessed motor-task fMRI. Data were originally acquired on a 3T Siemens Skyra using EPI (72 slices; in-plane FOV = 208 mm × 180 mm; 2.0-mm isotropic voxels; TR = 720 ms; TE = 33.1 ms; flip angle = 52°; BW = 2290 Hz/Px; multiband acceleration factor = 8). The HCP minimal preprocessing (Glasser et al., 2013) includes motion correction, EPI distortion correction, registration to MNI standard space, intensity normalization. We then performed additional preprocessing on the minimally preprocessed data, including spatial smoothing (FWHM = 4 mm), high-pass temporal filtering (cutoff = 200 s), and autocorrelation correction using standard FEAT procedures. Following (Barch et al., 2013), task-evoked activation maps were estimated using a 3-level FEAT analysis in FSL. The resulting activation maps were used to define task-specific ROIs for each movement type, and their spatial patterns were verified to correspond to canonical motor-task activations reported in (Barch et al., 2013), including the motor cortex, the somatosensory cortex, and the cerebellum.

Timeseries of each movement block were then extracted from these ROIs. In each scan, the last 12 seconds of two fixation blocks were also extracted from these ROIs to match the 12-second duration of the two corresponding movement blocks for each type of movement. The final dataset for each type of movement included four sets of movement timeseries per participant, with the corresponding fixation timeseries serving as the null condition for comparison.

#### 2.2.2. Simultaneously recorded local field potential (LFP)-fMRI data in rats

To assess the specificity of channel capacity and its cross-modal consistency, we analyzed concurrent LFP-fMRI recordings in anesthetized rats, using the resting brain as a negative-control setting in which no systematic hemispheric asymmetry is expected. In addition, to characterize the neural activity underlying the BOLD signal, we decomposed the LFP into band-limited power (BLP) time series across six frequency bands, estimated channel capacity from each BLP time series, and examined its correspondence with BOLD-derived channel capacity.

The data were acquired previously (Pan et al., 2011; Pan et al., 2013) through animal experiments approved by the Emory University Institutional Animal Care and Use Committee in compliance with NIH guidelines. Data preprocessing and quality control followed (Zhang et al., 2020). Briefly, male Sprague-Dawley rats underwent bilateral microelectrode implantation in the forelimb region of the primary somatosensory cortex (S1) and were subsequently scanned under subcutaneous dexmedetomidine anesthesia during simultaneous LFP-fMRI acquisition (Pan et al., 2011). LFP was recorded with 1000× amplification, band-pass filtered at 0.1-5000 Hz, notch filtered at 60 Hz, and digitized at 12 kHz. fMRI data were acquired on a 9.4 T small-animal system using EPI (TR = 500 ms, TE = 15 ms; 1020 volumes including 20 dummy scans), yielding approximately 10 min of data per scan.

MRI radiofrequency pulses induced brief artifacts in the LFP signal, which were used for temporal alignment between modalities and for segmentation into TR-locked epochs. Artifact templates were estimated by averaging across epochs and then subtracted, and residual artifact segments were replaced by linear interpolation. Epochs corresponding to dummy fMRI volumes were excluded. The remaining LFP signal was low-pass filtered at 100 Hz and downsampled to 500 Hz. Band-limited power was computed using 1-s sliding windows centered on each fMRI volume for six frequency bands: delta (1-4 Hz), theta (4-8 Hz), alpha (8-12 Hz), low beta (12-25 Hz), high beta (25-40 Hz), and gamma (40-100 Hz). Broadband power was calculated as the sum of power across all six bands. Under dexmedetomidine, the infraslow range was defined as 0.01-0.25 Hz, following prior simultaneous LFP-fMRI work in rats (Pan et al., 2013). Infraslow power (0.01-0.25 Hz) was additionally computed by band-pass filtering the full LFP signal and then applying a 1-s moving average to the squared signal.

For fMRI, a brain mask was generated from the first volume (active contour) and dilated by 2 voxels. Preprocessing in SPM12 included motion correction, spatial smoothing (Gaussian kernel, FWHM = 2.8 voxels; 0.84 mm), global signal regression, linear drift regression, and band-pass filtering (0.01–0.25 Hz). Two fMRI ROIs were selected as in prior work, corresponding to regions showing maximal cross-correlation (lag = 2.5 s) with LFP signals from the left and right hemispheres, respectively (Pan et al., 2011; Pan et al., 2013; Zhang et al., 2020). Following established quality-control (QC) procedures (Zhang et al., 2020), including QC on LFP residual noise, head motion, DVARS, bilateral correlation, and cross-modality correlation, 22 scans from 10 rats were retained for analysis. Additional acquisition details of image acquisition and preprocessing procedures were described in (Pan et al., 2011; Zhang et al., 2020).

#### 2.2.3. Simultaneously recorded wide field fluorescent calcium imaging (Ca^2+^)-fMRI data in mice

We further leveraged a simultaneously acquired wide-field calcium imaging–fMRI dataset in mice (hereafter denoted as Ca^2+^-fMRI for simplicity) to capture brain-wide temporal dynamics in both neuronal activity, reflected by intracellular calcium transients, and BOLD signals. These extended concurrent recordings provide a multimodal resource for examining time-varying neuronal and BOLD dynamics at the network level.

The data were acquired previously (Lake et al., 2020; Mandino, Horien, et al., 2025) through animal experiments approved by Yale Institutional Animal Care and Use Committee in compliance with NIH guidelines. Nine C57BL/6J mice expressing GCaMP6f in excitatory pyramidal neurons (Slc17a7/CaMKII/tet-GCaMP6f) underwent skull-thinning procedure to render the skull optically transparent, yet still intact, before implanting a head-plate to provide stable optical access to the dorsal cortical surface, with a fluorescent bead affixed for motion correction. Each animal participated in three imaging sessions separated by at least one week; each session comprised multiple scans of Ca^2+^-fMRI, with structural MRI scans interleaved for registration. During imaging, mice were maintained under light isoflurane anesthesia (0.5–0.75%, 70/30 medical air/O_2_).

Resting-state fMRI was acquired on an 11.7 T Bruker MRI scanner using EPI (TR = 1.0 s, TE = 9.1 ms; 0.4-mm isotropic voxels; 28 slices). Ca^2+^ imaging covered the dorsal cortex (field of view approximately 14 mm × 14 mm; 25 µm × 25 µm spatial resolution) with interleaved GCaMP-sensitive (470 nm) and control (395 nm) illumination at 20 Hz. The control channel was regressed from the sensitive channel to reduce background fluorescence, yielding an effective sampling rate of 10 Hz (Mandino, Horien, et al., 2025). Ca^2+^ images were motion-censored based on fMRI framewise displacement traces (threshold = 0.075 mm), spatially smoothed (σ = 0.1 mm), downsampled by a factor of 2, regressed against the control channel, and converted to 𝛥𝐹/𝐹₀. The resulting data were then registered to a CCFv3-based common template, as shown in Fig. 2 of (Mandino, Horien, et al., 2025).

**Fig. 2.**
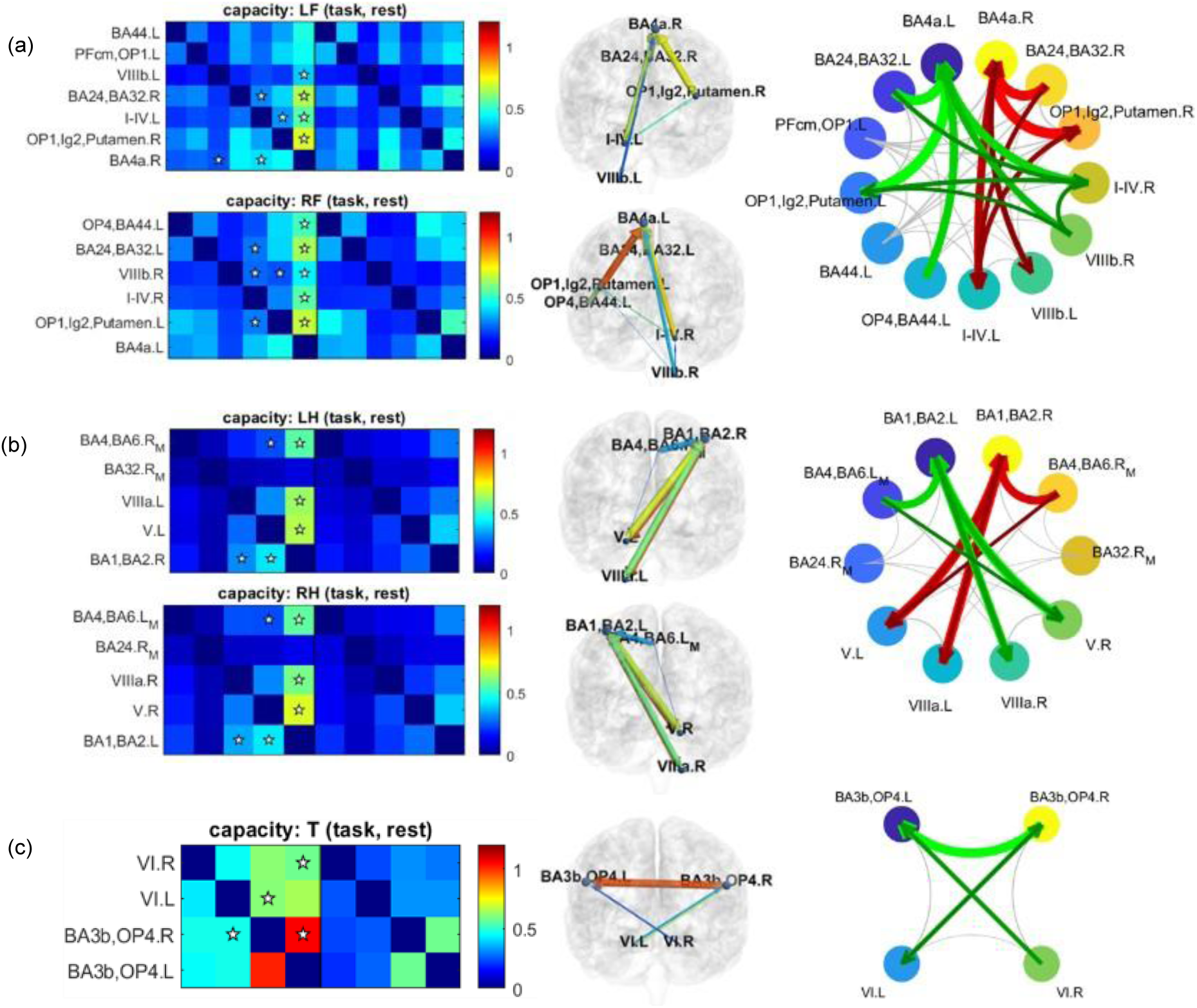
Motion-evoked directed connectivity among task-specific ROIs. Directed connectivity during foot movement (a), hand movement (b), and tongue movement (c) was identified using channel capacity at the 98% confidence level. For each movement type, significant motion-evoked connections are marked with ☆ in the connectivity matrices (left), rendered on the brain cortex using BrainNet Viewer ((Xia et al., 2013); middle), and summarized as connectograms (right). In the cortical renderings, arrow size reflects channel capacity and arrow color reflects p-corr strength. In (a) and (b), left- and right-sided motion-evoked connections are shown together in the connectograms, with left-sided connections in red and right-sided connections in green. In (c), tongue-evoked connections are shown in green. In the connectograms, foot and hand movements show largely mirror-symmetric left- and right-sided patterns consistent with contralateral sensorimotor organization, whereas tongue movement shows a more bilateral pattern.

For subsequent frequency-specific analyses, the Ca^2+^ signals were band-pass filtered into the infraslow (0.02-0.1 Hz) and delta (1-4 Hz) ranges. These bands are consistent with prior wide-field calcium imaging studies in mice under isoflurane anesthesia (Mitra et al., 2018). For clarity, we refer to the preprocessed signal without additional band-pass filtering as the broadband Ca^2+^ signal, in contrast to the infraslow and delta components.

fMRI preprocessing was performed using RABIES (Desrosiers-Grégoire et al., 2024), included framewise rigid head motion and mapping to Allen Atlas CCfv3 space via session-specific structural registration. In Allen space, fMRI nuisance regression included motion parameters, scrubbing high-motion frames (FD > 0.075 mm), spatial smoothing (FWHM = 0.8 mm), band-pass filtering (0.008–0.2 Hz), and global signal regression. Ca^2+^ data were registered to Allen space using the structural MRI described in (Mandino, Horien, et al., 2025). Timeseries of both Ca^2+^ and BOLD signals were then extracted from a subset of 60 ROIs from the Allen Atlas, spanning the cortical regions covered by the Ca^2+^ imaging FOV, for subsequent analysis.

### 2.3. Evaluation of Channel Capacity on Functional Neuroimaging Datasets

#### 2.3.1. Validation of channel capacity sensitivity in task-fMRI data

As described in Table 1, HCP motor task fMRI data was used to assess the sensitivity of channel-capacity–based directed connectivity to task-evoked changes. We expect that motion-evoked directed connectivity among task-relevant ROIs, quantified by channel capacity, is stronger during motor-task trials than during rest trials.

For each movement type, we pooled all task trials across subjects to form the task sample set and pooled the corresponding rest trials to form the rest sample set. Within each trial, we extracted ROI BOLD time series and estimated directed connectivity for each ordered ROI pair by fitting the trial-wise model described in Section 2.2 and computing the resulting channel capacity. Because each trial is short (12 s or 16 TRs), model parameters and channel capacity were estimated over the entire trial using a single window (i.e., no sliding-window segmentation). The corrected Akaike information criterion (AICc) was used to select the trial-specific model order 𝑘_𝑖_within each trial, which is expected to outperform AIC and BIC in small-sample settings (Hurvich & Tsai, 1989).

We defined motion-evoked connections as directed ROI pairs that exhibit (i) low between-subject variability in task-evoked channel capacity (CV < 30%) and (ii) a significant task-related increase relative to rest, assessed with within-subject, right-tailed paired t-tests (task vs. rest) and FDR-corrected across ROI pairs (q < 0.02).

To benchmark the task sensitivity of channel capacity, we used pairwise Granger causality (GC), a standard and interpretable directed-connectivity method. The pairwise form of GC is defined at the ROI-to-ROI level, which makes it methodologically comparable to our framework and avoids introducing unfair advantages or disadvantages that can arise from multivariate model specification. In (Smith et al., 2011), pairwise GC implementations were retained/preferred over several higher-order alternatives within the GC family. For our dataset, we use the Granger B4 variant described in (Smith et al., 2011). Specifically, Granger causality matrices were computed separately for task and rest conditions by estimating pairwise F-statistics across all ROI pairs. The model order for each ROI pair was selected using an information criterion, consistent with our framework. For GC, however, the model order should remain small enough to avoid over-parameterizing the autoregressive model, which can reduce estimation stability and statistical efficiency in fMRI GC analyses. In our subsequent analyses, the maximum tolerable lag was set to 4 TRs. Significant directional influences were identified using pairwise t-tests across the ROI dimension, following the same procedure described above.

#### 2.3.2. Validation of channel capacity specificity in resting state LFP-fMRI data

In resting state, we do not expect a consistent population-level directional bias between homotopic interhemispheric pairs; thus, directed connectivity estimates should be approximately symmetric across hemispheres under the resting-state null (Moon et al., 2025). If channel capacity is a specific measure of directionality (i.e., with a low false-positive rate), then its estimates from the resting-state LFP-fMRI data should show significantly smaller directional asymmetry compared to its scan-to-scan variability. The directional asymmetry is defined as the difference in time-averaged connectivity between one connection direction and its opposite direction. The scan-to-scan variability is defined as the standard deviation of the time-averaged connectivity across all scans. For each scan, the time-averaged connectivity is calculated by averaging the connectivity measure across all sliding windows within that scan.

We formulate the null hypothesis 𝐻_0_ : median directional asymmetry ≥ scan-to-scan variability and apply a left-tailed Wilcoxon signed-rank test. A p-value < 2%, leads us to reject 𝐻_0_, supporting the interpretation that directional asymmetry is insignificant compared to baseline scan-to-scan fluctuations, consistent with a low false-positive tendency in inferred directionality.

Following prior sliding-window correlation analysis on this dataset (Thompson et al., 2013b), we used 50-second sliding windows with a 1 TR step. Following (Xu et al., 2017, 2021, 2026), we used the BIC to select the model order 𝑘_𝑖_ within each window. We performed the above-mentioned statistical test on channel capacity across all modalities (fMRI and six LFP BLPs) of this dataset, pooling sliding windows across all scans to form the sample set.

Notably, our prior work has demonstrated low directional asymmetry (i.e., high specificity) for SWpC strength and connection duration in resting-state data (Xu et al., 2026). Here, we extend this validation framework to the newly introduced channel-capacity measure, providing a direct and comparable assessment of its directionality specificity under a resting-state symmetry expectation.

#### 2.3.3. Multimodal evaluation of resting-state dynamic directed FC estimated by channel capacity

To evaluate whether channel capacity provides a cross-modally consistent and neurobiologically meaningful measure of directed FC in the resting brain, we leveraged concurrently acquired LFP and Ca^2+^ imaging as neural readouts alongside their matched fMRI data. LFP provides high temporal resolution and was decomposed into distinct band-limited power (BLP) signals, whereas wide-field Ca^2+^ imaging provides cortex-wide coverage with high spatial resolution. For each modality, we estimated channel capacity as a measure of time-varying information transfer, yielding a sequence of windowed capacity estimates.

##### Time-resolved correspondence and resting-state dynamics

To assess whether channel capacity captures shared time-varying dynamics across modalities, we analyzed the resting-state LFP-fMRI dataset by computing windowed channel capacity estimates for fMRI and for each BLP signal using the same sliding-window parameters (50 s windows, 1 s step). For each directed ROI pair (or predefined directed connection) we obtained one capacity estimate per window, yielding paired time series 𝐶^fMRI^(𝑤) and 𝐶^BLP^(𝑤) across windows 𝑤 . We quantified cross-modal time-resolved correspondence by computing the correlation between these paired window-wise capacity time series, and estimated uncertainty using bootstrap resampling of windows (resampling windows with replacement within scan and aggregating across scans).

##### Brain-wide, time-averaged cross-modal correspondence

For the Ca^2+^-fMRI data, we computed scan-level, time-averaged channel-capacity matrices for each modality by averaging the windowed channel-capacity estimates across all sliding windows within a scan (45 s windows, 1 s step). We quantified Ca^2+^–fMRI correspondence by computing the correlation between the connectivity matrices derived from fMRI and from band-limited Ca^2+^ signals in either the infraslow (0.02–0.1 Hz) or the delta (1–4 Hz) band. Uncertainty was estimated using bootstrap resampling across scans.

##### Brain-state transitions and cross-modal state correspondence

To characterize temporally alternating resting-state organization in a brain-wide setting, we analyzed the resting-state Ca^2+^-fMRI dataset by computing a directed channel-capacity connectivity matrix for each sliding window in each modality. We focused on fMRI and the infraslow WF-Ca^2+^ band (0.02–0.1 Hz) due to their similar spectral content. Each window’s directed capacity matrix was treated as one sample; matrices from all windows (45 s windows, 1 s step) across all scans were pooled, vectorized over directed off-diagonal entries, and clustered using *k*-means to identify discrete connectivity states and brain-state transitions over time. We evaluated solutions for *k* = 1, …,10 (using six random initializations for stability) and used the elbow method to determine the optimal *k*. Specifically, the optimal number of clusters was selected using the elbow method based on the within-cluster sum of squares (WCSS) curve (Vergara et al., 2020). To compare the cluster centroids derived from fMRI data versus Ca^2+^ data, we fixed the labeling of the fMRI centroids and permuted the Ca^2+^ centroids. The permutation that maximized the total correlation across the *k* pairs of centroids was selected to align and relabel the Ca^2+^ centroids. To benchmark the clustering results obtained with channel capacity, we repeated the same analysis using SWC. The resulting fMRI centroids from channel capacity and SWC were then aligned using the same procedure.

## 3. Results

### 3.1. Channel Capacity Sensitively Captures Motor-evoked Information Transfer

HCP motor-task fMRI data provide a natural testbed for evaluating whether channel capacity can detect biologically meaningful task-evoked directionality by contrasting movement and rest periods. As shown in Fig. 2, the motion-evoked directed connections recovered by channel capacity were consistent with the laterality of the movements and with known sensorimotor physiology.

For hand and foot movements, the dominant directed interactions were from the cerebellum to the contralateral somatomotor cortex, consistent with established sensorimotor circuitry (Fieblinger, 2021; Rizzolatti & Luppino, 2001). For example, left foot movement involved VIIIb.L→BA4a.R, right foot movement involved VIIIb.R→BA4a.L, left hand movement involved VIIIa.L→BA1/BA2.R, and right hand movement involved VIIIa.R→BA1/BA2.L. Channel capacity also identified directed interactions from OP1-Lg2-putamen to BA4a during both left and right foot movements.

Additional interactions consistent with motor-sensory integration were also identified, such as BA4/BA6 → BA1/BA2 (Todorov, 2004). For tongue movement, channel capacity detected bilateral cerebellar-to-tongue-area interactions (VI.L→BA3b/OP4.R; VI.R→ BA3b/OP4.L), consistent with the cerebellum’s role in fine motor control and complex orofacial movements (Manto et al., 2011; Sasegbon & Hamdy, 2023). Together, these findings indicate that channel capacity recovers canonical sensorimotor lateralization and somatotopic organization in the motor network.

We next benchmarked this motion sensitivity against pairwise Granger causality (GC-B4 in (Smith et al., 2011)). Using the same thresholds (CV < 30% and q < 0.02), GC did not detect any significant connections. Detecting at least one significant connection for each movement type required relaxing the threshold to q < 0.24, corresponding to a 76% confidence level, substantially lower than the 98% confidence level used for channel capacity. Under this relaxed threshold, the GC-derived connections (Fig. S2) did not consistently reflect movement laterality, and several physiologically relevant pathways were missed. For example, during left foot movement, GC did not identify VIIIb.L→ BA4a.R or OP1-Lg2-putamen.R→BA4a.R, and during left hand movement it did not detect BA4/BA6.RM → BA1/BA2.R.

Together, these results show that channel capacity detects task-evoked directed interactions with high sensitivity while preserving biologically meaningful lateralization and circuit structure.

### 3.2. Channel Capacity Reliably Detects Directionality with High Specificity

We then assessed the specificity of channel capacity by applying sliding-window analysis to bilateral resting-state LFP-fMRI data in rats. Because the two hemispheres are expected to exhibit ongoing interhemispheric coupling in the resting state, but not a stable, systematic directional bias favoring one hemisphere over the other, this dataset provides a negative-control setting for evaluating false-positive directionality.

The Wilcoxon signed-rank test (Fig. 3) shows that the median directional asymmetry is significantly lower than the scan-to-scan variability (p < 1e-3) using channel capacity estimates. This indicates that channel capacity does not detect directionality between ROI pairs when no such directionality is expected, thereby validating its specificity for directionality. Furthermore, this result holds consistently across the fMRI and each LFP BLP, demonstrating robust cross-modality reliability of channel capacity. The same calculations were also performed for p-corr strength and connection duration and validated their directional specificity as well (Fig. S3).

**Fig. 3.**
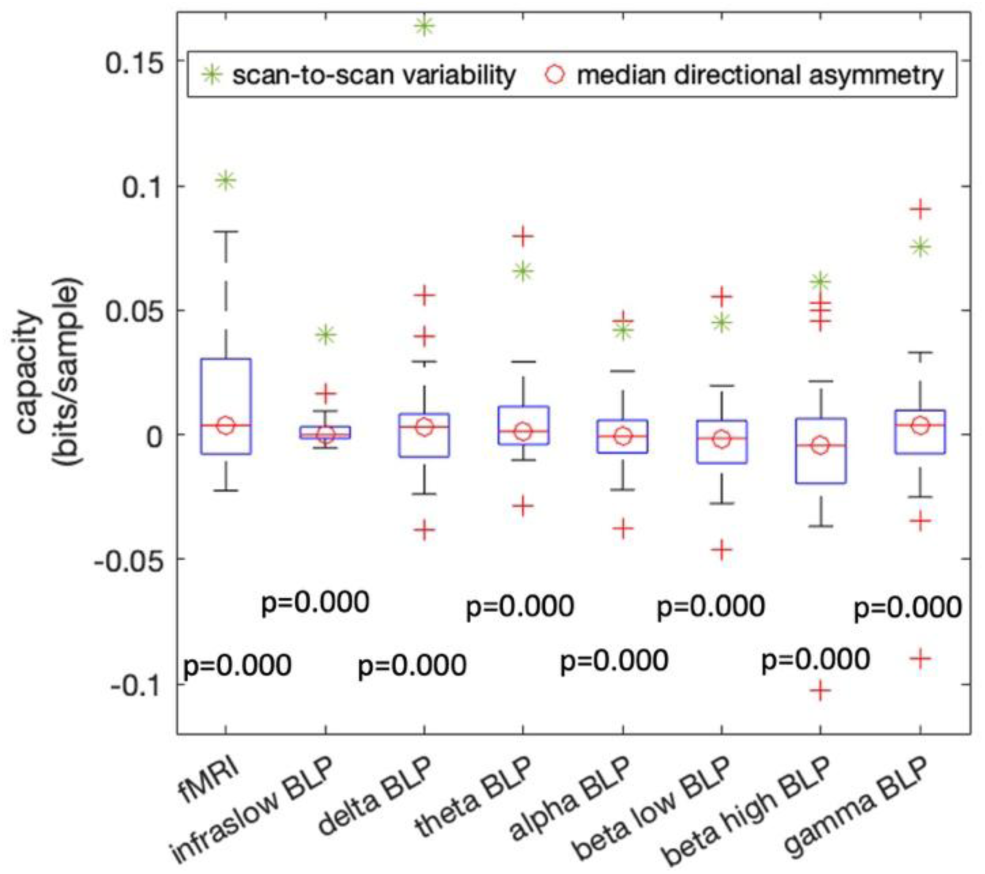
Directional asymmetry of channel capacity in the LFP-fMRI dataset. The p-values of the Wilcoxon signed-rank tests between directional asymmetry and scan-to-scan variability are shown below each boxplot.

### 3.3. Associations Between BOLD and Neuronal Directed FC Measured by Channel Capacity

Beyond establishing sensitivity and specificity, we next asked whether channel capacity captures directed interactions that are neurobiologically meaningful across modalities. We address this question in two complementary ways. First, we test whether BOLD-derived directed FC covaries with simultaneously measured neural signals in expected frequency ranges. Second, we ask whether channel capacity reveals recurring whole-brain resting-state patterns and whether those patterns show cross-modal correspondence between BOLD and infraslow neural signals. Together, these analyses assess whether channel capacity reflects not only directional structure, but also temporally organized neural dynamics.

#### 3.3.1. Associations in frequency-specific directed FC

This analysis asks whether channel capacity estimated from BOLD fMRI tracks directed connectivity estimated from concurrent neural recordings, and if so, in which frequency ranges. A positive cross-modal association would suggest that the BOLD-derived measure is not arbitrary, but reflects neural communication structure captured by the simultaneously recorded LFP or Ca^2+^ signals. Because these neural signals span multiple timescales, frequency-specific differences are also informative about which components of neural activity are most closely aligned with BOLD-derived directed FC.

Bootstrap correlations of time-resolved channel capacity sequences between BOLD fMRI and LFP BLPs are shown in Fig. 4a. No significant positive correlation is observed between BOLD and delta or alpha BLPs. The correlations of BOLD with theta BLP and with infraslow BLP are relatively strong but only borderline significant (p ≈ 0.05). Positive correlations between BOLD and high frequency BLPs (beta low, beta high, and gamma) are significant (p < 0.05) but systematically decrease from beta low to beta high and from beta high to gamma. In Fig. S4a, we repeated the same analysis for sliding-window p-corr strength, correlation, and connection duration. Except for connection duration, which showed near-zero correlation, the other connectivity measures produced correlation distributions similar to channel capacity for each BLP. This suggests that channel capacity can serve as a measure of cross-modality association, producing results comparable to those obtained with correlation-based measures.

**Fig. 4.**
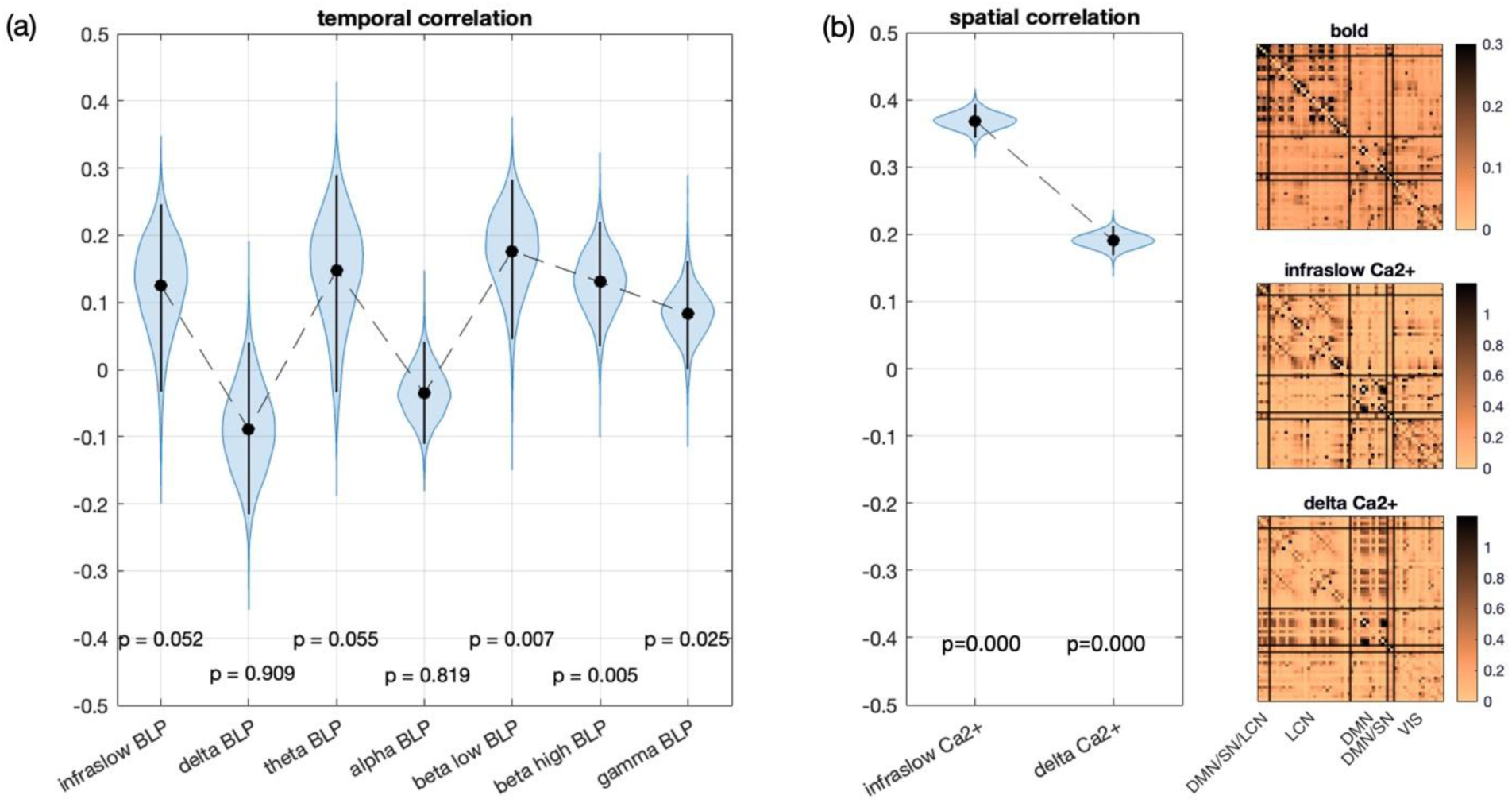
Cross-modal comparison of BOLD-derived and neural (LFP/Ca^2+^-derived) channel capacity. (a) Bootstrapped temporal correlations between concurrently acquired LFP- and BOLD-derived channel-capacity time series for each LFP band-limited power (BLP) band. (b) Ca^2+^-fMRI results: bootstrapped spatial correlations between concurrently acquired Ca^2+^- and BOLD-derived time-averaged channel-capacity matrices for the infraslow and delta components of the Ca^2+^ signals (left), together with the corresponding time-averaged channel-capacity matrices for BOLD, infraslow Ca^2+^, and delta Ca^2+^ (right). P-values indicate whether the bootstrapped correlation is greater than zero.

Bootstrap correlations of time-averaged channel capacity matrices between BOLD fMRI and Ca^2+^ signals in infraslow band and in delta band are shown in Fig. 4b. Both Ca^2+^ frequency bands show significant positive correlations with the BOLD signal (p < 1e-4 for both); however, the correlation in the delta band is much weaker than that in the infraslow band, consistent with the result previously reported (Mandino et al., 2025; Mitra et al., 2018; O’Connor et al., 2022). The same comparison can also be observed in the time-averaged connectivity matrices averaged across all scans (Fig. 4c). Both the BOLD and infraslow Ca^2+^ matrices show strong connectivity in the lateral cortical network (LCN), while the delta band Ca^2+^ matrix shows strong connectivity in the default mode network (DMN) and its coupling with LCN. The other connectivity measures we tested (Fig. S4b) showed weaker effects: the difference in p-corr was not significant, the difference in correlation was reversed, and connection duration yielded weak correlations in both frequency bands. This suggests that channel capacity can more effectively distinguish informative neural communications from unrelated signals due to the temporal memory incorporated in the connectivity model.

Taken together, these results show that cross-modal agreement is selective rather than uniform across neural frequencies and modalities. In the LFP-fMRI data, BOLD-derived channel capacity aligns most consistently with higher-frequency BLPs and only marginally with theta and infraslow fluctuations, whereas in the Ca^2+^-fMRI data the strongest spatial correspondence is observed for the infraslow component.

#### 3.3.2. Associations in time-resolved directed FC and brain states

Given the strong spatial correspondence between infraslow neural and BOLD channel-capacity matrices observed in Fig. 4b, we next examined whether window-resolved channel-capacity matrices revealed recurring brain-wide patterns in both modalities.

Resting-state brain dynamics are commonly assessed using brain states identified through clustering. We applied k-means clustering to window-resolved channel-capacity estimates from the resting-state Ca^2+^-fMRI dataset. The elbow method identified 𝑘 = 4 as the optimal number of clusters in both BOLD and Ca^2+^ datasets (as shown in Fig. S5).

The fMRI cluster centroids for 𝑘 = 4 are shown in Fig. 5a (top row). State 1 is the weakest-connectivity state. State 2 exhibits the strongest overall connectivity, characterized by strong within- and between-network connectivity involving the default mode network (DMN) and the lateral cortical network (LCN). State 3 is characterized by prominent lateralized connectivity in the LCN and visual (VIS) network and by stronger coupling between these networks and others. State 4 shows prominent connectivity involving the DMN, the LCN, and the salience network (SN).

**Fig. 5.**
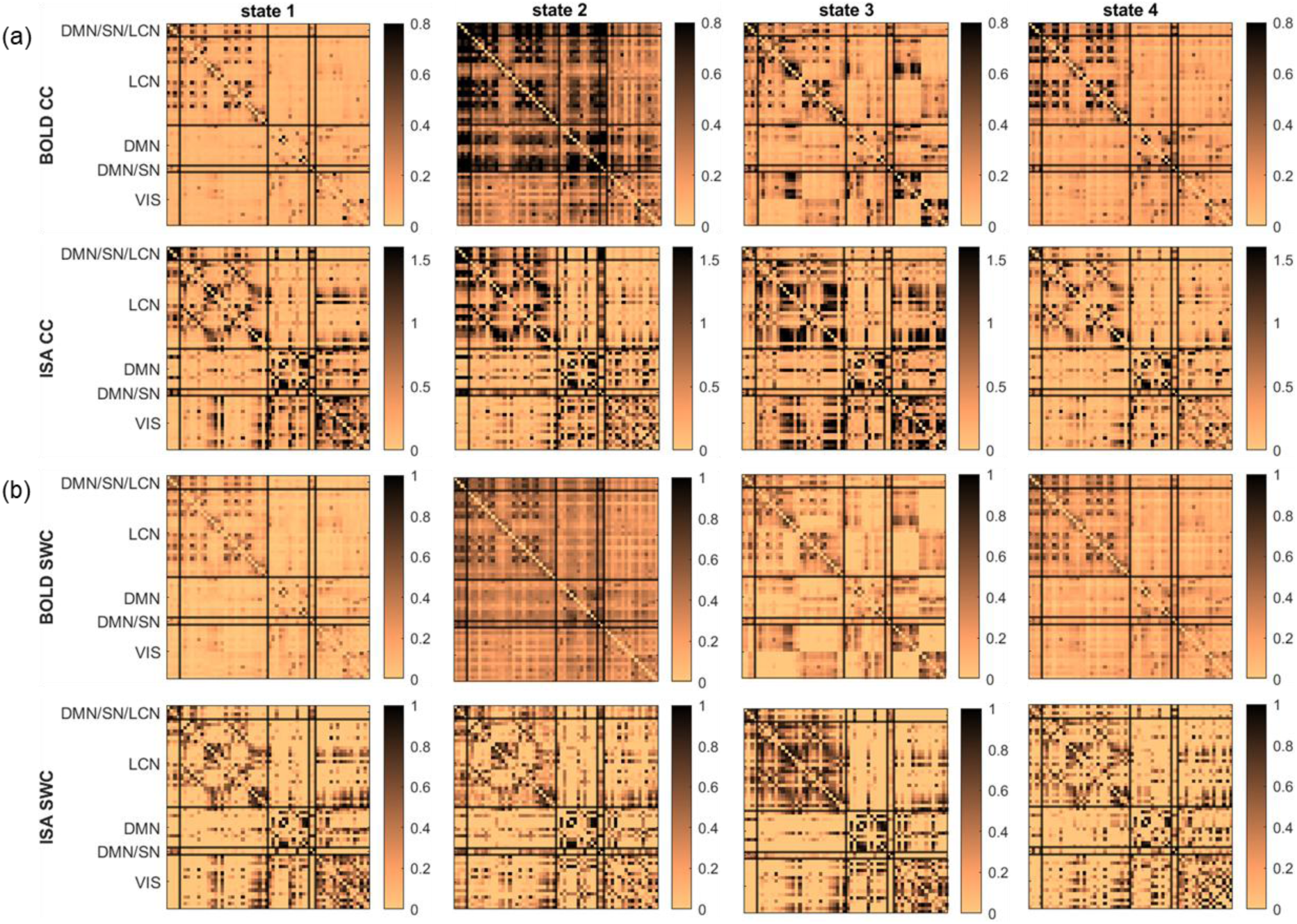
Channel capacity reveals recurring brain-wide resting-state patterns with partial cross-modal correspondence between BOLD and infraslow neural (ISA) signals. (a) Channel-capacity (CC)-based state centroids obtained by k-means clustering of window-resolved connectivity matrices pooled across scans. The top row shows BOLD fMRI and the second row shows ISA signals; columns correspond to the four recurring states. ISA state labels were aligned to the BOLD-derived states by maximizing centroid similarity across modalities. (b) Sliding-window-correlation (SWC)-based state centroids shown for comparison. The third row shows BOLD SWC states and the fourth row shows ISA SWC states. In all matrices, ROIs are grouped by functional network (DMN/SN/LCN, LCN, DMN, DMN/SN, and VIS), and color bars indicate connectivity magnitude.

The infraslow Ca^2+^ cluster centroids for 𝑘 = 4 are shown in Fig. 5a (second row). The connectivity patterns in the Ca^2+^ centroids are noticeably different from those observed in fMRI, and the distinctions between states are more subtle. Nevertheless, several structured patterns are apparent. In State 1, the LCN exhibits selective connectivity and coupling with the VIS network. In State 2, the LCN shows strong coupling with the motor area (DMN/SN/LCN) and the cingulate area (DMN/SN). In State 3, the VIS network is highly connected and strongly coupled with the LCN. State 4 is characterized by overall weaker connectivity compared with the other states.

For comparison, the cluster centroids obtained using SWC are shown in Fig. 5b. These centroids exhibit substantial similarity to those derived from the channel-capacity analysis.

Table S1 shows that centroids obtained with channel capacity and SWC are highly correlated across both fMRI (≥ 0.90) and Ca^2+^ (≥ 0.64). The correlation of the state-alternation time courses between channel capacity and SWC is 0.66 ± 0.21 in fMRI and 0.31 ± 0.44 in Ca^2+^ (Fig. S7). Additionally, Table S2 shows that the correspondence between fMRI and Ca^2+^ centroids is higher for channel capacity than for SWC.

Together, these results show that channel capacity identifies recurring, structured brain-wide states in resting data, with partial cross-modal correspondence between BOLD and infraslow neural signals. This suggests that the directed interactions captured by channel capacity reflect organized large-scale brain dynamics rather than unstructured temporal variation.

### 3.4. Relation between Channel Capacity and Other Connectivity Measures

To better understand what channel capacity captures relative to existing measures, we compared its time course with those of correlation, p-corr and duration of information transfer. In a typical scan (Fig. 6), channel capacity shows temporal variation in both magnitude and directionality that is broadly consistent with p-corr and connection duration, while appearing more sensitive than p-corr to changes in directional magnitude associated with connection duration. Numerical comparisons summarized in Table 2 confirm that channel capacity is strongly correlated with p-corr. However, this relationship depends on connection duration. As shown in Fig. 7, the 𝑅^2^between channel capacity and p-corr decreases as connection duration increases. When the connection duration 𝑘_𝑖_ = 1 TR, corresponding to no memory effect in ROI interactions, channel capacity and p-corr follow a deterministic and monotonically increasing relationship. When 𝑘_𝑖_ = 2 TRs, their relationship begins to show dispersion, and this dispersion increases further as 𝑘_𝑖_ increases. Together, these results indicate that channel capacity is closely related to p-corr, but is not reducible to it, because it captures additional features of connectivity when information flow between ROIs is temporally sustained.

**Fig. 6.**
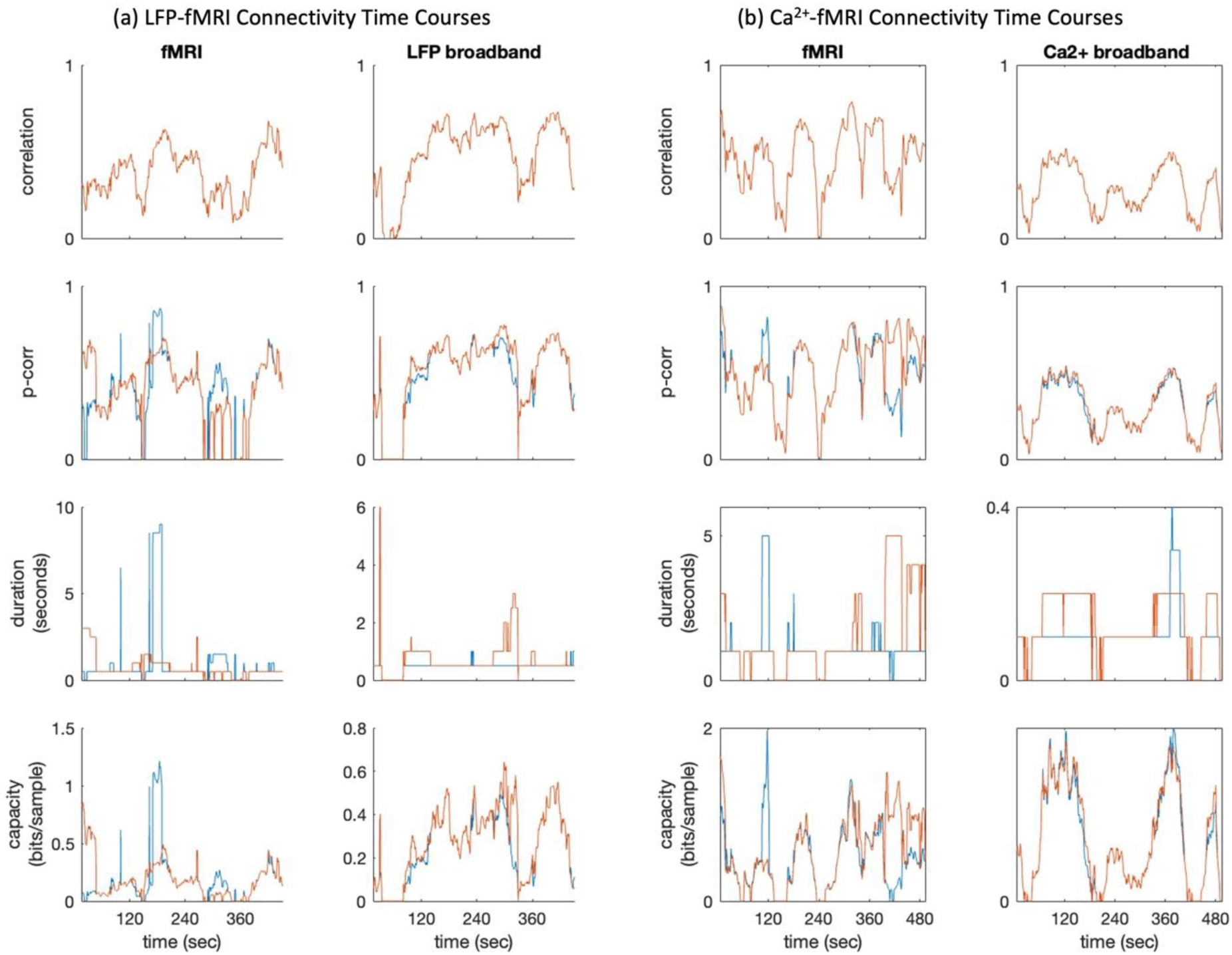
**Connectivity time courses during a representative scan from the multimodal resting-state datasets**: (a) the LFP-fMRI dataset and (b) the Ca^2+^-fMRI dataset. Blue traces indicate connectivity from S1L to S1R, and red traces indicate connectivity from S1R to S1L. For correlation, the two traces overlap because correlation is a nondirectional measure.

**Fig. 7.**
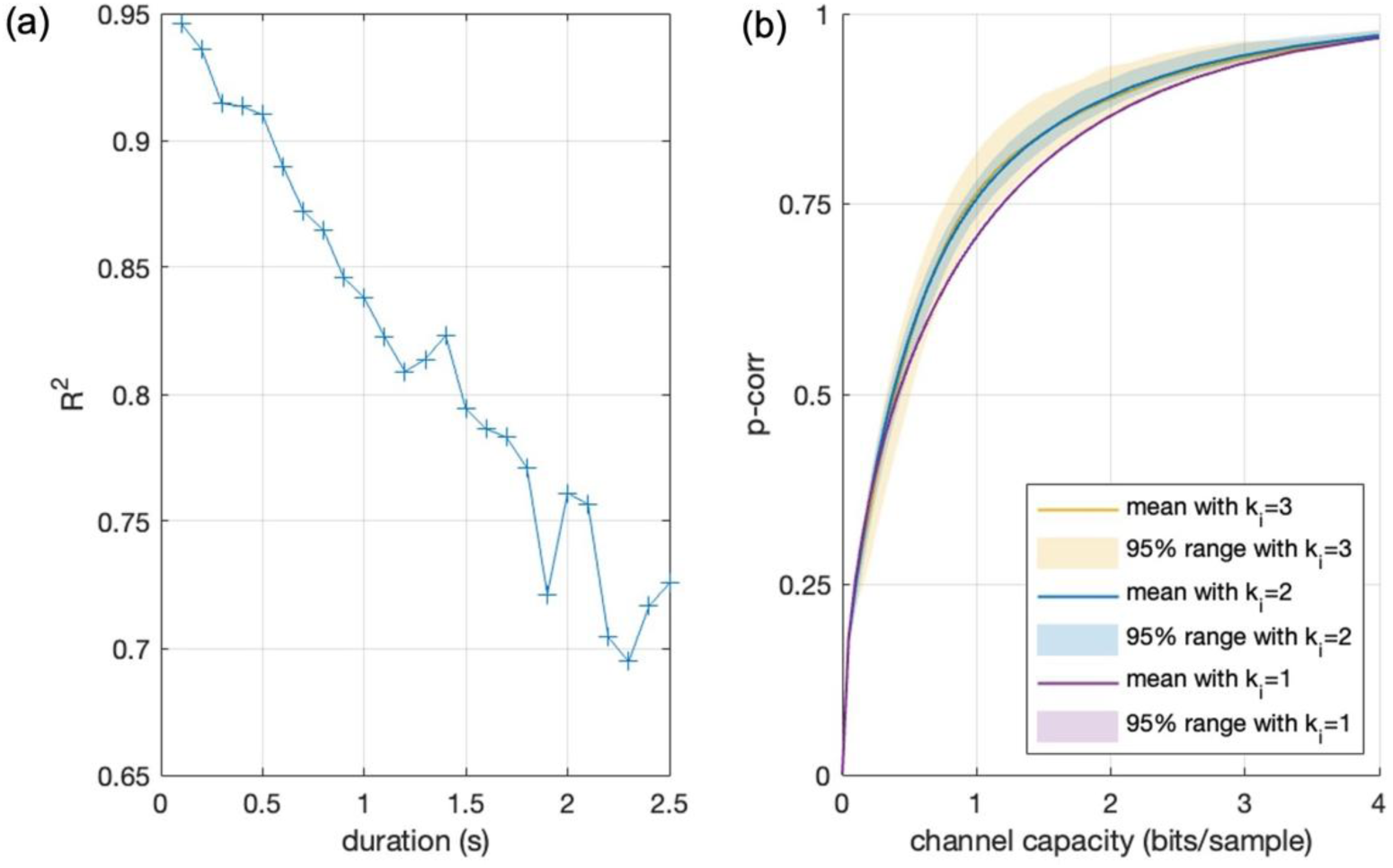
Channel capacity increasingly diverges from p-corr as connection duration increases. (a) 𝑅^2^ between channel capacity and p-corr as a function of connection duration 𝑘_𝑖_, estimated using all connectivity samples from the Ca^2+^-fMRI data across windows, scans, and directed ROI pairs. (b) Connectivity samples plotted in the channel-capacity–p-corr plane and color-coded by connection duration 𝑘_𝑖_ (up to 3 TRs).

**Table 2.** Correlations between the time course of channel capacity and the time courses of other connectivity measures for S1L-to-S1R connectivity in each dataset.

| Correlation between: | fMRI | LFP broadband | fMRI | Ca<sup>2+</sup> imaging |
| --- | --- | --- | --- | --- |
| capacity, p-corr | 0.94±0.04 | 0.96±0.02 | 0.96±0.03 | 0.98±0.03 |
| capacity, correlation | 0.69±0.26 | 0.85±0.14 | 0.89±0.14 | 0.95±0.11 |
| capacity, duration | 0.70±0.25 | 0.52±0.26 | 0.56±0.32 | 0.48±0.37 |

## 4. Discussion

This paper introduced channel capacity as a model-based, information-theoretic measure of dynamic effective connectivity. Under a finite impulse response (FIR) channel model with additive white Gaussian noise (AWGN), channel capacity quantifies the maximum achievable information-transfer rate supported by the inferred directed interaction, providing a physically interpretable complement to conventional connectivity-strength measures. Because it can be estimated within the same framework as p-corr and its sliding-window extension, channel capacity is readily scalable to large ROI sets and long-duration signals while also supporting time-resolved analysis.

Across multimodal datasets in humans and rodents, channel capacity demonstrated several desirable properties for dynamic effective-connectivity analysis, including sensitivity to physiologically meaningful task-evoked interactions, specificity against spurious directionality in resting state, sensitivity to brain-wide resting-state organization, and meaningful correspondence with concurrent neural and BOLD measurements. Importantly, these validations were established across complementary datasets, each probing a distinct aspect of the method.

We first evaluated channel capacity in HCP motor-task fMRI, where the experimental design and known somatotopic organization provide a clear benchmark for sensitivity. In this dataset, channel capacity reliably identified task-evoked directed interactions, recovering canonical sensorimotor pathways, including cerebellum-to-contralateral somatomotor cortex effects, as well as somatotopically organized patterns across foot, hand, and tongue movements (Fieblinger, 2021; Mandino, Horien, et al., 2025; Rizzolatti & Luppino, 2001; Sasegbon & Hamdy, 2023). These findings indicate that channel capacity is sensitive to biologically meaningful directional interactions in a setting where expected task modulation is well established. This is an important validation, because sensitivity in directed-connectivity analysis is often difficult to demonstrate without either invasive perturbation or strong prior physiological expectations. Here, the motor-task paradigm provided a practical benchmark showing that channel capacity can recover interpretable and reproducible directed interaction patterns in human fMRI. In this benchmark, channel capacity also showed higher practical sensitivity than pairwise Granger causality, recovering physiologically coherent task-evoked interactions under stricter statistical thresholds.

We next extended the evaluation to rodent resting-state multimodal datasets, where concurrent neural measurements enabled more stringent assessment of specificity and cross-modal validity. In the resting-state rat LFP-fMRI dataset, channel capacity exhibited negligible directional asymmetry relative to scan-to-scan variability across both fMRI and LFP band-limited power signals. Because no strong systematic hemispheric directionality is expected in this negative-control setting, this result supports a low false-positive rate in directionality estimation. This property is particularly important for dynamic effective-connectivity methods, since apparent asymmetries can easily arise from noise, model mismatch, or preprocessing-related artifacts. The rat LFP-fMRI results therefore provide evidence that channel capacity is not only sensitive, but also appropriately conservative when persistent directionality is not expected.

Beyond specificity, the rodent multimodal datasets also enabled assessment of whether channel capacity captures neurobiologically meaningful cross-modal structure. In the resting-state mouse Ca^2+^-fMRI dataset, channel capacity revealed cross-modal correspondence at both the brain-wide, time-averaged level and the dynamic state level. At the brain-wide, time-averaged level, channel-capacity matrices derived from fMRI were positively correlated with those derived from Ca^2+^ signals, with stronger correspondence observed for the infraslow Ca^2+^ band than for the delta band, consistent with the slow temporal structure of resting-state BOLD fluctuations. The stronger correspondence for infraslow Ca^2+^ is consistent with (Mitra et al., 2018), who showed that spontaneous infraslow calcium activity has unique large-scale spatiotemporal dynamics and is reflected in hemodynamic/BOLD signals. At the dynamic level, clustering of windowed channel-capacity matrices revealed recurring resting-state connectivity states and state transitions over time. Within each modality, these state patterns were highly similar to those obtained with SWC, supporting the robustness of the recovered large-scale dynamic organization. Across modalities, however, the correspondence between infraslow Ca^2+^- and BOLD-derived states was partial rather than one-to-one, consistent with prior simultaneous Ca^2+^-fMRI studies showing both common and divergent patterns of cortical functional organization (Vafaii et al., 2024), while extending those findings to the level of time-varying brain-state organization.

Taken together, the LFP-fMRI and Ca^2+^-fMRI findings suggest that channel capacity can serve not only as a directed-connectivity measure within a single modality, but also as a useful framework for assessing cross-modal correspondence between hemodynamic and neural signals. In the LFP-fMRI data, channel-capacity fluctuations in BOLD were associated with corresponding fluctuations in LFP band-limited power signals. In the Ca^2+^-fMRI data, channel capacity captured both time-averaged and state-level correspondence between BOLD and Ca^2+^-derived connectivity structure. These results support the interpretation that channel capacity reflects biologically meaningful aspects of neural communication underlying BOLD dynamics.

Channel capacity was also not redundant with existing connectivity measures. Relative to pairwise Granger causality, it showed higher practical sensitivity in the motor-task benchmark, recovering more physiologically coherent task-evoked interactions under stricter statistical thresholds. Relative to SWC, it recovered similar large-scale state structure within modality while showing stronger cross-modal correspondence between BOLD and infraslow Ca^2+^ signals. Relative to p-corr and related measures, it remained strongly associated in simple cases but diverged as connection duration increased, indicating sensitivity to temporally sustained aspects of directed interaction.

Channel capacity quantifies the maximum rate at which information can be transmitted through a communication channel under specified signal and noise constraints. Applied to brain imaging, this idea provides an interpretable way to quantify effective connectivity: rather than asking only whether two brain signals are statistically related, it asks how much directed information transfer the fitted interaction can support. Unlike commonly used information-theoretic dependence measures such as transfer entropy (Vicente et al., 2010), that quantify statistical dependencies between signals, channel capacity directly characterizes the communication channel itself, thereby providing a description of the underlying substrate supporting directed interaction under the fitted model. Importantly, channel-capacity estimation explicitly incorporates noise statistics, an essential feature for functional neuroimaging data, where signal-to-noise ratio is often low and spatially heterogeneous across ROIs. In this sense, channel capacity offers a mechanistic lens that aligns naturally with the concept of directed information flow in brain networks. At the same time, because it can be estimated within the sliding-window framework, it remains computationally practical and well suited for time-resolved analysis.

These properties make channel capacity a promising tool for functional neuroimaging for assessing dynamic effective connectivity. More broadly, channel capacity provides a bridge between the practical feasibility of dynamic FC approaches and the directionality of EC approaches, while retaining a clear physical interpretation. Because it can be estimated within a sliding-window framework, it is also computationally practical and well suited for time-resolved analysis.

Nevertheless, several limitations should be noted. First, channel capacity estimates depend on model assumptions. This paper presented a relatively simple model to establish proof-of-concept, including linear FIR with Gaussian noise. In future work, we will investigate more realistic channel models by incorporating nonlinearity, feedback mechanisms, and non-Gaussian noise to improve the robustness and explanatory power of the proposed approach. Second, this paper adopted a pairwise channel model without explicitly accounting for multivariate confounding and network-level structure. Extending the current framework using conditional or partialized multivariate formulations will be an important next step.

In summary, time-resolved channel capacity provides a physiologically grounded, dynamic, and directed measure for effective connectivity, that complements existing FC and EC approaches. By first demonstrating sensitivity in human motor-task fMRI and then establishing specificity and cross-modal neural relevance in rodent multimodal resting-state datasets, this work provides proof-of-concept that channel capacity can recover meaningful directed information-flow organization in functional neuroimaging. These findings motivate further development of information-theoretic approaches for assessing large-scale dynamic effective connectivity in functional neuroimaging.

## Code Availability

The code used in this study is available at https://github.com/inspirelab-site/swcc.

## Author Contributions

Jianan Jian: data curation; data preprocessing; methodology; formal analysis; writing, original draft.

Benjamin Li: data preprocessing; formal analysis; writing, review. Nurahmed Multezem: initial analysis on mouse Ca^2+^-fMRI dataset.

Francesca Mandino: data acquisition; data curation; data preprocessing on mouse Ca^2+^-fMRI dataset; writing, review.

Evelyn Lake: data acquisition; data curation; data preprocessing on mouse Ca^2+^-fMRI dataset; writing, review.

Nan Xu: Conceptualization; methodology; data preprocessing; data curation; formal analysis; writing, review and editing; funding acquisition; and supervision.

## Acknowledgement

This research is supported by NIH R00NS123113. Data were provided in part by the Human Connectome Project, WU-Minn Consortium (Principal Investigators: David Van Essen and Kamil Ugurbil; 1U54MH091657) funded by the 16 NIH Institutes and Centers that support the NIH Blueprint for Neuroscience Research; and by the McDonnell Center for Systems Neuroscience at Washington University. We also thank Dr. Keilholz group in providing the LFP-fMRI data in rats.

## Supplementary Material

Supplementary material associated with this article is provided.

## Footnotes

1 We selected 𝑁 = 512 for a sufficient frequency resolution for the spectral approximation.

