## Supplementary Figures and Tables for "Channel Capacity for Time-Resolved Effective Connectivity in Functional Neuroimaging"

### Supplementary Materials

(LF)

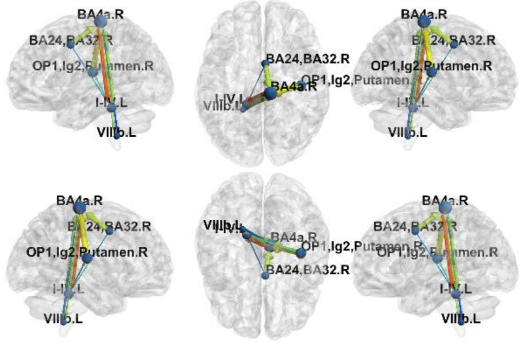


(RF)

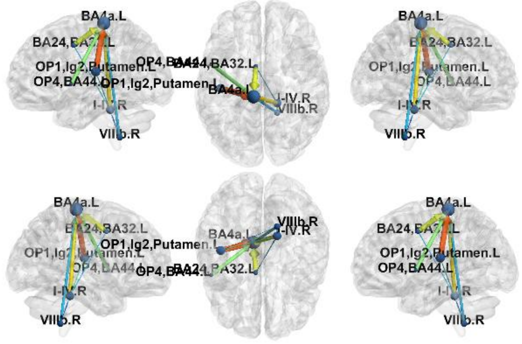


(LH)

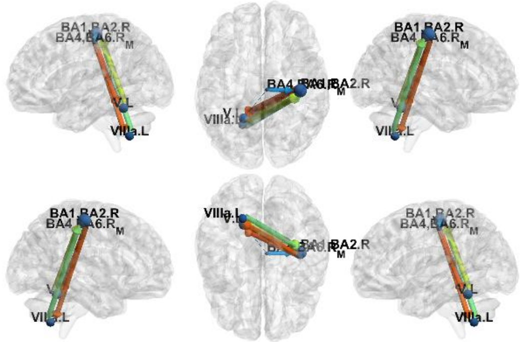


(RH)

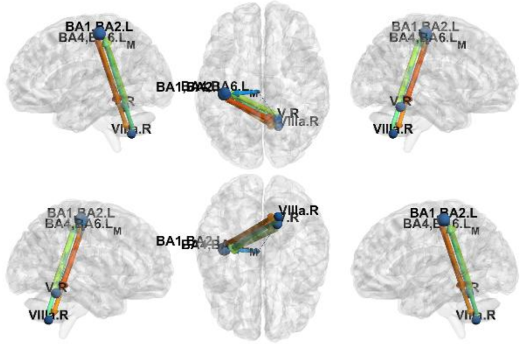


(T)

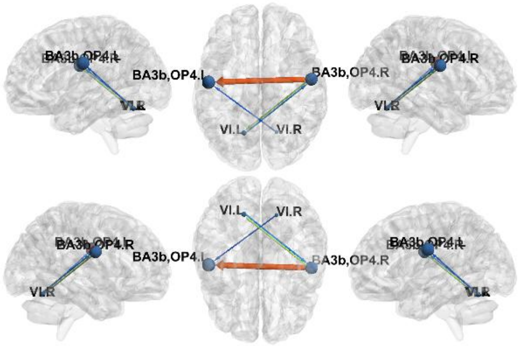


**Fig. S1 *Sagittal, axial, and coronal views of the motion-evoked connections.*** Directed connectivity in each movement type (LF, left foot; RF, right foot; LH, left hand; RH, right hand; T, tongue) was identified among task-specific ROIs using channel capacity at the 98% confidence level. Arrow size reflects channel capacity and arrow color reflects p-corr strength.

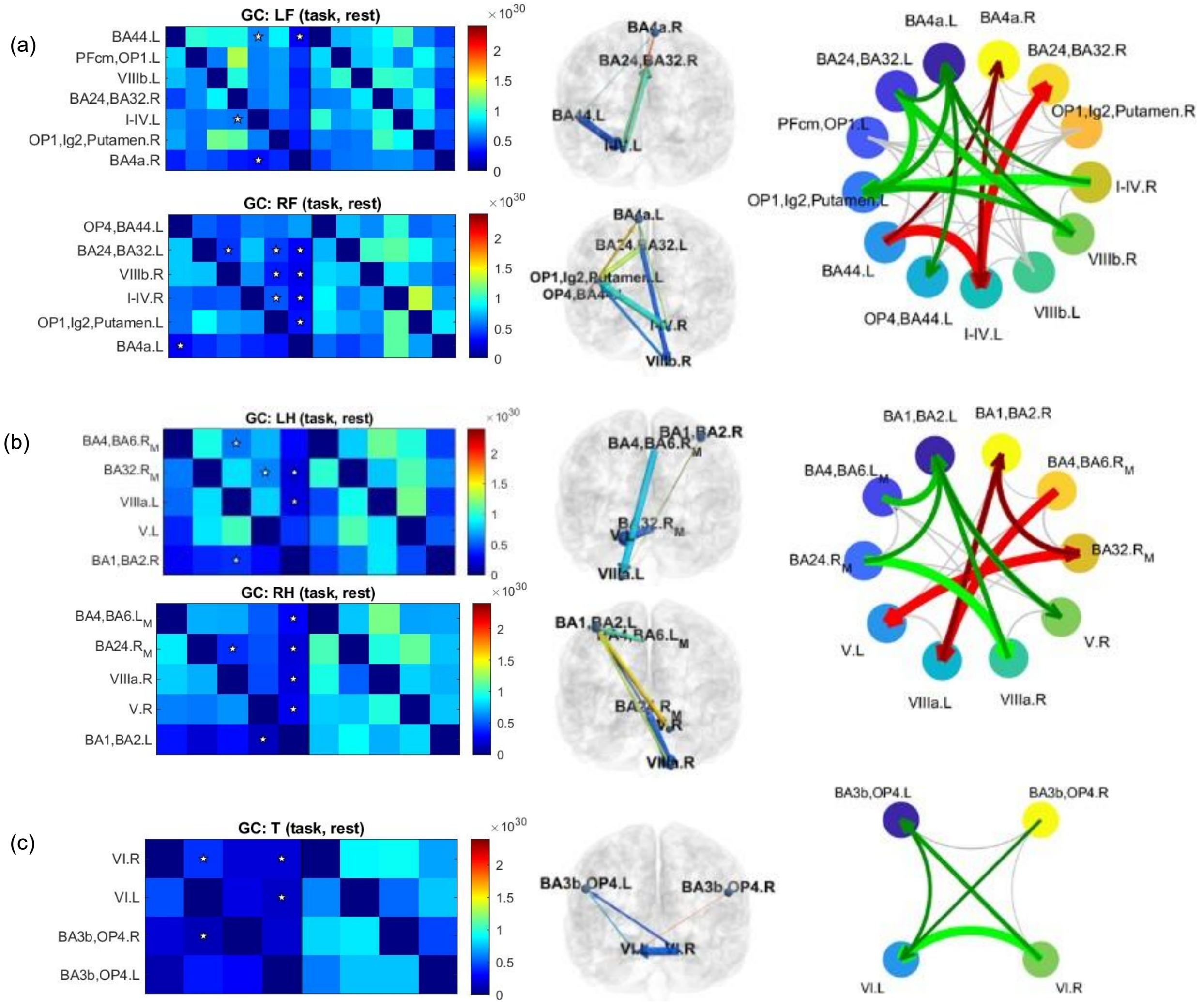


**Fig. S2 Motion-evoked directed connectivity identified using Granger causality.** Directed connectivity during foot movement (a), hand movement (b), and tongue movement (c) was identified among task-specific ROIs using channel capacity at the 76% confidence level. For each movement type, significant motion-evoked connections are marked with ☆ in the connectivity matrices (left), rendered on the brain cortex using BrainNet Viewer ((Xia et al., 2013); middle), and summarized as connectograms (right). In the cortical renderings, arrow size reflects Granger causality and arrow color reflects p-corr strength. In (a) and (b), left- and right-sided motion-evoked connections are shown together in the connectograms, with left-sided connections in red and right-sided connections in green. In (c), tongue-evoked connections are shown in green.

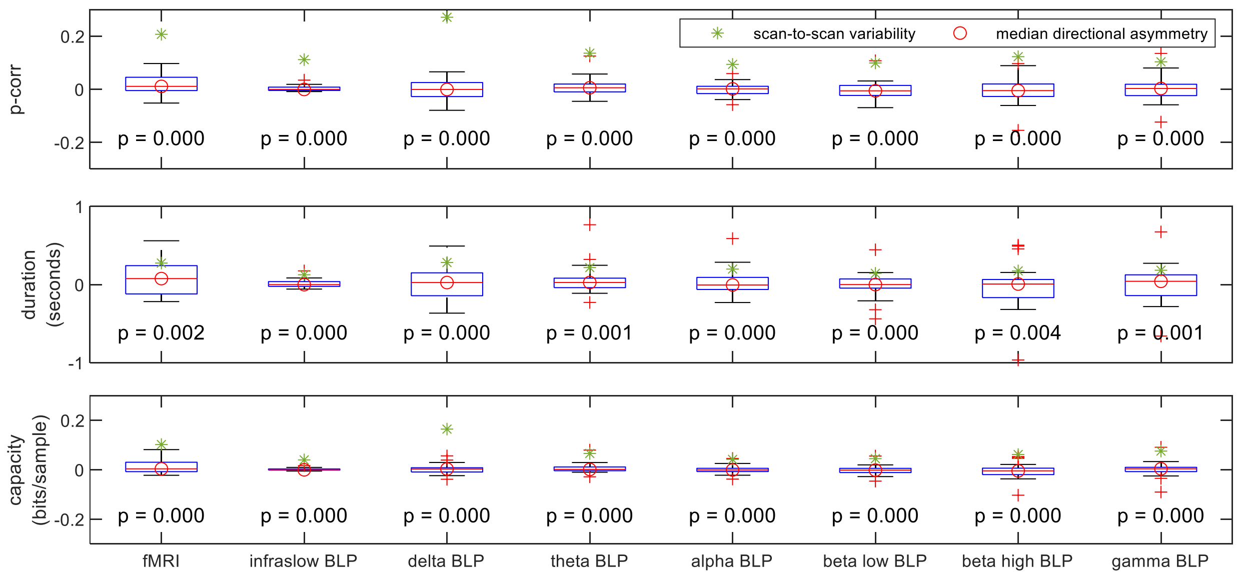


**Fig. S3 Directional asymmetry of p-corr (top), duration (middle), and channel capacity (bottom) in the LFP-fMRI dataset.** The p-values of the Wilcoxon signed-rank tests between directional asymmetry and scan-to-scan variability are shown below each boxplot.

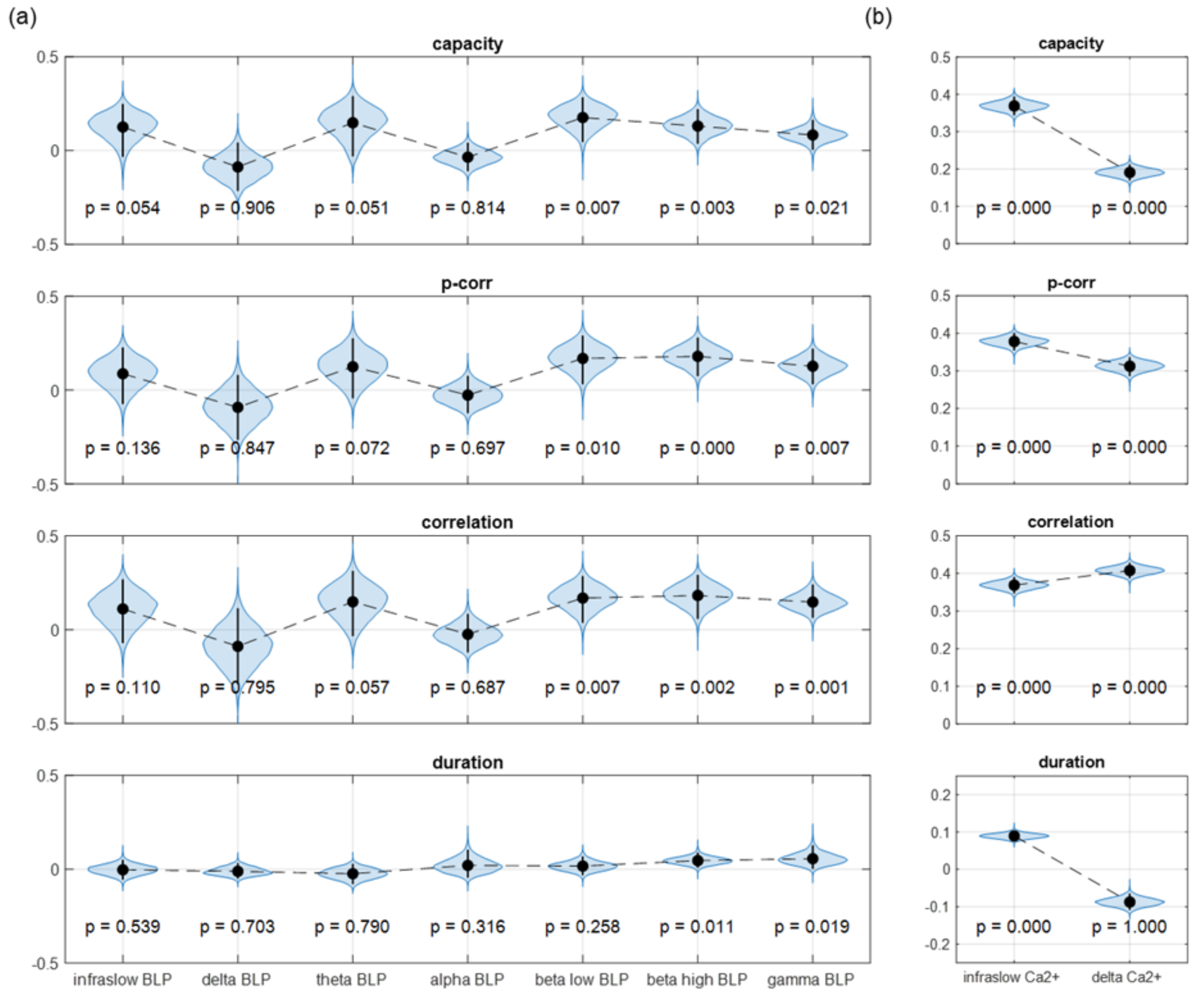


**Fig. S4 Cross-modal comparison of BOLD-derived and neural (LFP/Ca2+-derived) connectivity measures (channel capacity, p-corr, correlation, duration).** (a) Bootstrapped temporal correlations between concurrently acquired LFP- and BOLD-derived connectivity time series for each LFP band-limited power (BLP) band. (b) $Ca^{2+}$-fMRI results: bootstrapped spatial correlations between concurrently acquired $Ca^{2+}$- and BOLD-derived time-averaged connectivity matrices for the infraslow and delta components of the $Ca^{2+}$ signals. P-values indicate whether the bootstrapped correlation is significantly greater than zero.

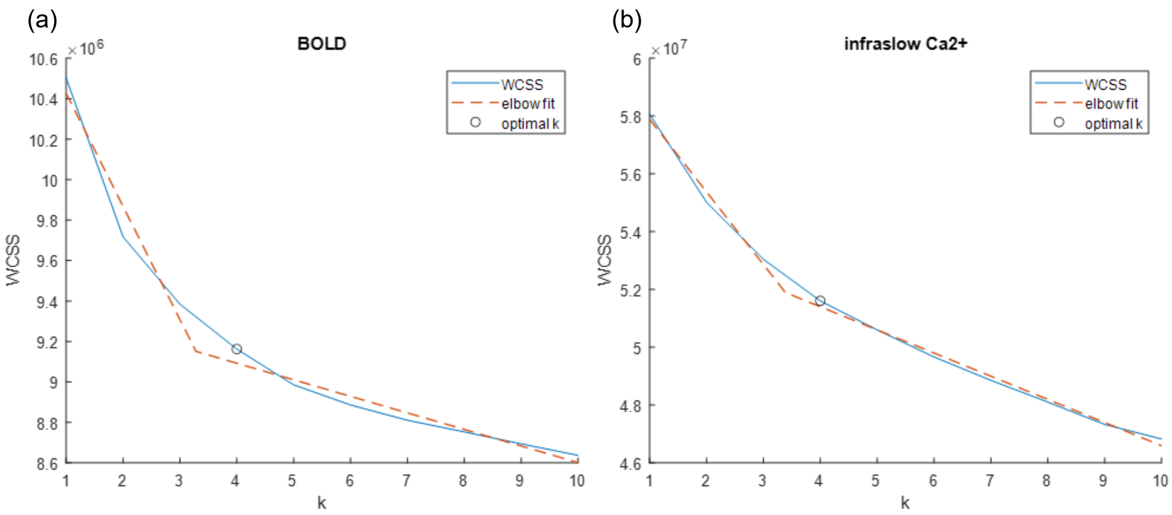


**Fig. S5 Elbow method for determining the number of clusters in resting-state data,** applied to (a) BOLD signals and (b) infraslow $Ca^{2+}$ signals. The solid blue line shows the within-cluster sum of squares (WCSS) as a function of cluster number k, and the dashed red line indicates the two-segment linear fit. The optimal k was chosen as the smallest integer greater than the breakpoint.

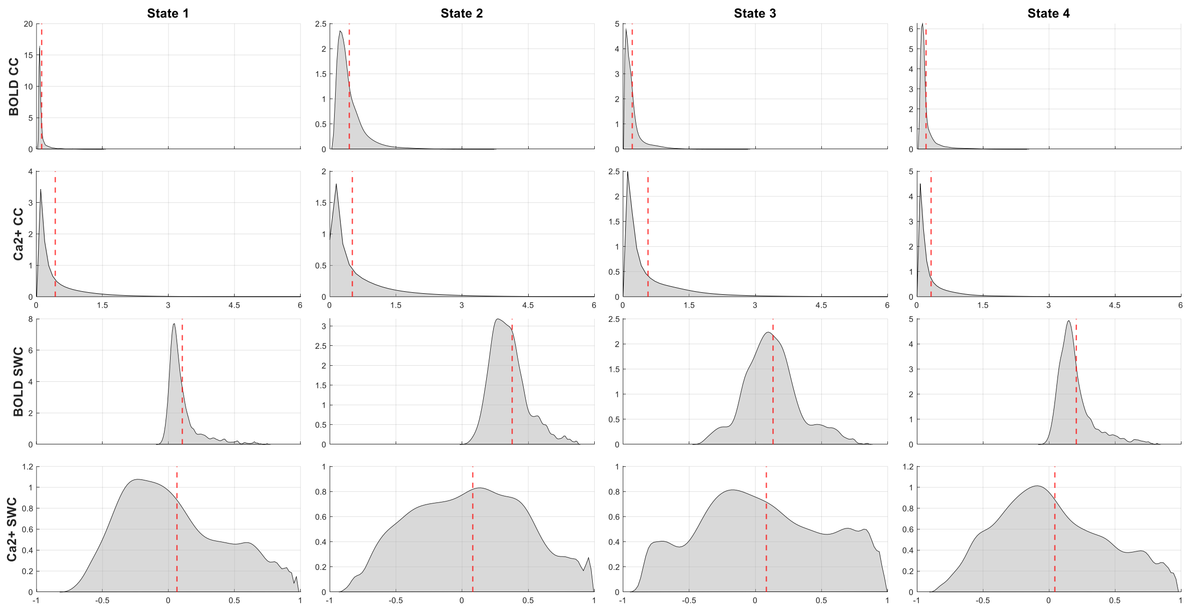


**Fig. S6 Probability distributions of connectivity values in each of the four states.** First row: channel capacity (CC) in the BOLD fMRI data. Second row: channel capacity (CC) in the infraslow band $Ca^{2+}$ data. Third row: sliding-window correlation (SWC) in the BOLD fMRI data. Fourth row: sliding-window correlation (SWC) the infraslow band $Ca^{2+}$ data. Dotted red lines mark the means.

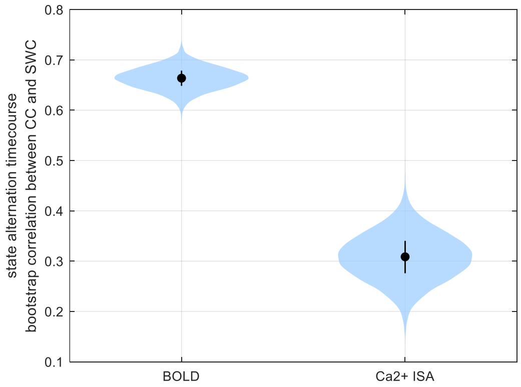


**Fig. S7 Bootstrap correlation of the state alternation time course between channel capacity and SWC.**

| | State 1 | State 2 | State 3 | State 4 |
|---|---|---|---|---|
| fMRI centroids | 0.90 | 0.96 | 0.94 | 0.97 |
| $Ca^{2+}$ centroids | 0.64 | 0.67 | 0.71 | 0.78 |

**Table S1. Correlation between channel capacity centroid and SWC centroid for each state.**

| | State 1 | State 2 | State 3 | State 4 |
|---|---|---|---|---|
| Capacity | 0.56 | 0.54 | 0.53 | 0.65 |
| SWC | 0.35 | 0.44 | 0.37 | 0.60 |

**Table S2. Correlation between fMRI centroid and $Ca^{2+}$ centroid for each state using either Capacity or sliding-window correlation (SWC) connectivity measure.**
